# Learning-based screening of hematologic disorders using quantitative phase imaging of individual red blood cells

**DOI:** 10.1101/091983

**Authors:** Geon Kim, YoungJu Jo, Hyungjoo Cho, Hyun-seok Min, YongKeun Park

**Affiliations:** Department of Physics, Korea Advanced Institute of Science and Technology (KAIST), Daejeon 34141, Republic of Korea; KAIST Institute for Health Science and Technology, KAIST, Daejeon 34141, Republic of Korea; Tomocube, Inc., Daejeon 34051, Republic of Korea

## Abstract

We present a rapid and label-free method for hematologic screening for diseases and syndromes, utilizing quantitative phase imaging (QPI) and machine learning. We aim to establish an efficient blood examination framework that does not suffer from the drawbacks of conventional blood assays, which are incapable of profiling single cells or using labeling procedures. Our method involves the synergistic employment of QPI and machine learning. The high-dimensional refractive index information arising from the QPI-based profiling of single red blood cells is processed to screen for diseases and syndromes using machine learning, which can utilize high-dimensional data beyond the human level. Accurate screening for iron-deficiency anemia, reticulocytosis, hereditary spherocytosis, and diabetes mellitus is demonstrated (>99% accuracy) using the proposed method. Furthermore, we highlight the synergy between QPI and machine learning in the proposed method by analyzing the performance of the method.

## 1. Introduction

The use of disease-induced alterations of individual red blood cells (RBCs) as diagnostic markers that may facilitate efficient screening for various diseases and syndromes has been widely reported (Baskurt et al. 1998; Bateman et al. 2017; Jung et al. 2016; Kim et al. 2014a; Koepke and Koepke 1986; Lee et al. 2017; Watanabe et al. 1994). Previous investigations have stated that the diverse properties of RBCs, such as cellular morphology and deformability, change depending on the pathophysiological conditions (Diez-Silva et al. 2010; Kim et al. 2012). The literature describes significant alterations of RBCs under a variety of pathophysiological conditions, including parasitic infection (Chandramohanadas et al. 2011; Kim et al. 2014a; Park et al. 2008b), inflammation (Baskurt et al. 1998; Bateman et al. 2017), and metabolic diseases (Kotisaari et al. 2002; Lee et al. 2017; Schmid-Schönbein and Volger 1976).

Unfortunately, conventional techniques for hematological examination offer limited access to the properties of individual red blood cells. An example of a conventional method is the complete blood count (CBC), which is the current gold standard technique for measuring RBC properties. In laboratory medicine, CBCs are routinely employed to obtain diagnostic parameters, including the mean corpuscular volume, mean corpuscular hemoglobin (Hb) content, and the ratio between different blood cell populations. Despite the well-established diagnostic protocols and usefulness of this approach in hematological studies (Higgins 2015; Higgins and Mahadevan 2010; Urrechaga et al. 2013; Weatherall 2011b), the massive *population-based* nature of CBC measurements and analyses suggest the clear limitation that this method is incapable of characterizing *individual* RBCs. Considering the increasing awareness of cellular heterogeneity at the single-cell level (Altschuler and Wu 2010; Wang and Bodovitz 2010), a new technique for profiling individual RBCs would play a major role in extending the potential of RBC-based disease screening (Weatherall 2011a).

Quantitative phase imaging (QPI), a recently emerging imaging technology based on laser interferometry, has been proven capable of rapidly characterizing individual blood cells (Kemper and von Bally 2008; Lee et al. 2013; Popescu 2011). Unlike conventional microscopes based on exogenous labeling (e.g., fluorescence microscopy or peripheral blood smear examination), QPI exploits the intracellular refractive index (RI) distribution as an endogenous imaging contrast for label-free imaging without exhaustive sample preparation. Holotomography, also known as three-dimensional (3D) QPI (Kim et al. 2014a; Wolf 1969), measures 3D RI tomograms of live cells and tissues, from which various parameters can be retrieved. For example, morphological (e.g., cellular volume) and biochemical (e.g., Hb content) properties of blood cells can be retrieved at the individual cell level (Kim et al. 2018; Kim et al. 2014b; Memmolo et al. 2014; Park et al. 2008a; Yoon et al. 2017). Furthermore, time-lapse two-dimensional (2D) QPI directly measures dynamic RBC membrane fluctuations, revealing the mechanical properties (e.g., deformability) of individual RBCs (Park et al. 2010a; Park et al. 2010b; Shaked et al. 2011). Combining 3D QPI with time-lapse 2D QPI for simultaneous access to all the aforementioned properties, or single-cell profiling of individual RBCs, has also been demonstrated (Kim et al. 2014c; Kim et al. 2014d).

However, diagnostic applications of QPI-based RBC profiling have been challenging, owing to the lack of analytical tools for fully exploiting the fruitful information. For the population-averaged parameters obtained through CBC, it has been possible for human experts to hand-craft diagnostic criteria. In contrast, single-cell profiling data is typically high-dimensional (i.e., it is difficult to visualize), and accompanies cell-to-cell variations that preclude rule-based diagnostic approaches.

Despite the difficulties, recent advances in machine learning suggest new opportunities in diagnostic applications (Cruz and Wishart 2007). The data-driven approach provides indispensable methods for analyzing high-dimensional and complicated data, as in single-cell profiling. In QPI, adopting machine learning has enabled rapid identification of bacterial species (Jo et al. 2015; Jo et al. 2017), sperm cell analysis (Mirsky et al. 2017), macrophage activation (Pavillon et al. 2018), and lymphocyte subtyping (Yoon et al. 2017), among a rapidly expanding range of applications (Jo et al. 2018). Recently, 2D QPI techniques have been combined with machine learning for the detection of abnormal morphologies in RBCs, such as malaria infections (Go et al. 2018) or sickle cell diseases (Javidi et al. 2018). No study to date has demonstrated the efficient handling of single-RBC profiling data incorporating all the morphological, chemical, and mechanical properties simultaneously.

Here, we present a systematic method to rapidly and efficiently screen for hematologic disorders using QPI and machine learning. In the proposed method, a neural network classifier predicts the pathophysiological conditions of individuals based on the simultaneously measured properties of individual RBCs. We show that single-cell profiling provides invaluable information in the context of high-accuracy screening for iron-deficiency anemia (IDA), reticulocytosis (RET), hereditary spherocytosis (HS), and diabetes mellitus (DM). Importantly, the key traits utilized by the classifier are consistent with the literature, even though the classifier was trained in a purely data-driven manner without any human knowledge. We also propose a strategy for overcoming and even exploiting the cell-to-cell variations for more accurate screening.

## 2. Materials and Methods

### 2.1. Profiling of individual RBCs

QPI images of RBCs were measured using common-path diffraction optical tomography (cDOT) (Kim et al. 2014c). 3D RI tomograms and time-lapse phase maps are measured for each RBC. The image acquisition process for 3D RI tomograms is visualized in Figs. 1A and B. Using multiple 2D optical field images of a sample obtained with various illumination angles, a 3D RI tomogram is reconstructed. The illumination angle is scanned using two two-axis galvanometric mirrors. One galvanometric mirror was placed before a sample at the conjugated plane, in order to control the illumination angle of the beam. The other was located after the sample, so that the optical axis of the transmitted beam remained unchanged. A common-path interferometer was employed to record an interferogram of a sample beam, from which the optical field is recovered (Debnath and Park 2011). The 3D RI tomogram of a specimen is reconstructed from the multiple 2D optical fields, using an optical diffraction tomography algorithm (Kim et al. 2013; Kim et al. 2016; Lauer 2002; Wolf 1969). The information in the spatial frequencies that are inaccessible owing to the limited numerical apertures of objective lenses is filled using an iterative non-negativity algorithm (Lim et al. 2015).

**Fig. 1.**
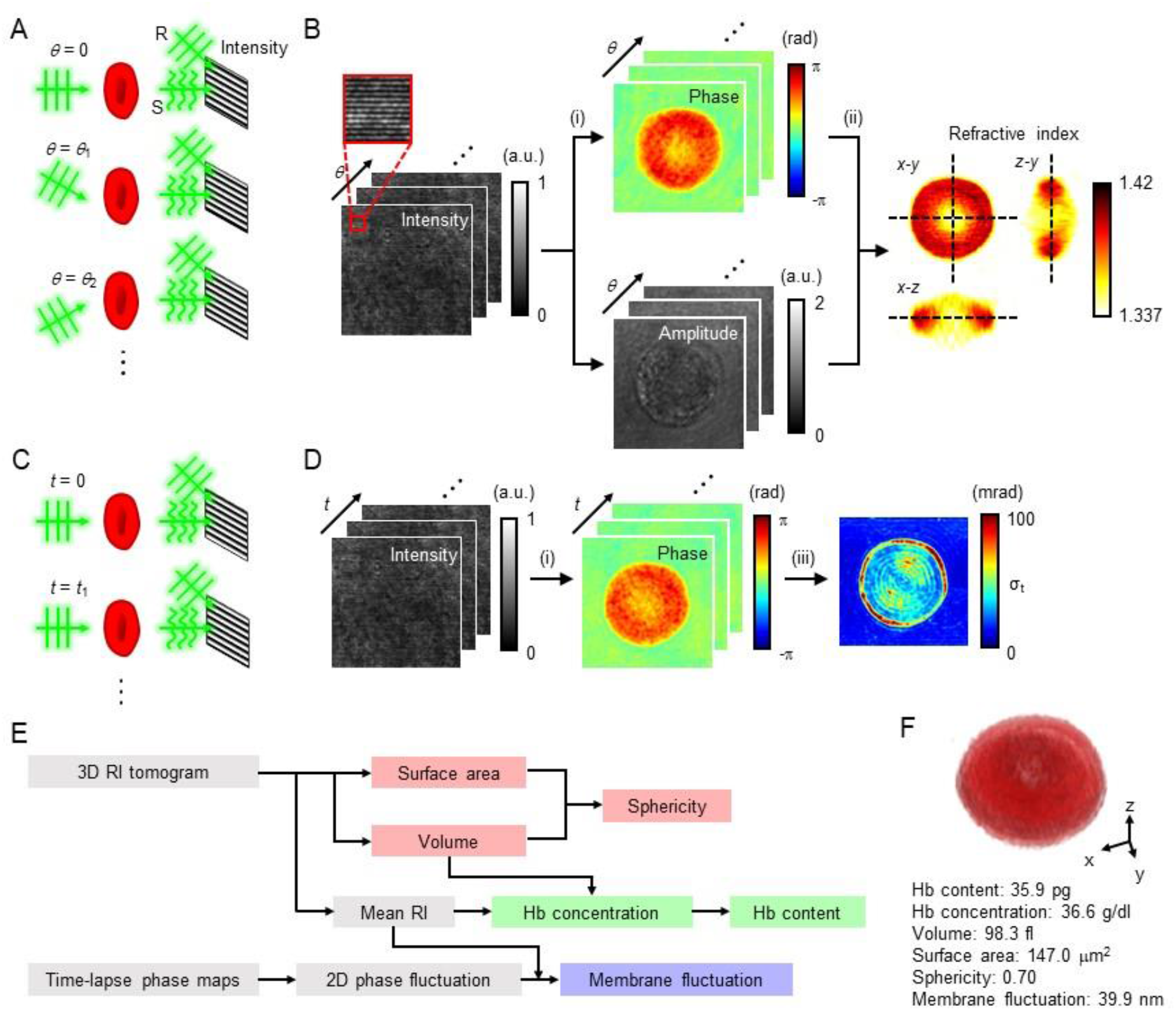
Profiling of single red blood cells using quantitative phase imaging. (A) The holographic measurement scheme for obtaining a three-dimensional (3D) refractive index (RI) image. S and R refer to the scattered wave and reference wave, respectively. (B) The process in which a 3D RI tomogram is reconstructed. (C) The holographic measurement scheme for obtaining time-lapse phase maps. (D) The process in which time-lapse phase maps are retrieved. (i), (ii), and (iii) refer to optical field retrieval, diffraction tomography, and temporal standard deviation, respectively. (E) A flow diagram illustrating the extraction of the cellular properties of a red blood cell (RBC) from the 3D RI tomogram and the 2D time-lapse phase images. (F) A 3D volume rendering corresponding to the RBC in (B) and the extracted properties.

The time-lapse 2D QPI acquisition process is illustrated in Figs. 1C and D. Time-lapse phase maps are retrieved from the measurements of 2D optical fields by considering the RI value of each RBC measured using 3D QPI. For each RBC, 256 2D QPI images were measured at the frame rate of 125 frames/s. The detailed procedure is described elsewhere (Kim et al. 2014d).

3D QPI is capable of profiling single RBCs by providing quantitative cellular features and high-resolution images in 3D. Six cellular properties, including volumes, surface areas, sphericities, Hb contents, Hb concentrations, and membrane fluctuations, were quantified from the QPI images using prior knowledge concerning RBCs (Lee et al. 2017). The volume and surface area of an RBC can be directly obtained from a 3D RI tomogram. The sphericity, a dimensionless index ranging from 0 to 1, represents the sphere-likeness of a given 3D object. This is calculated as the ratio of surface areas between a sphere with the identical volume and the given 3D object. The Hb concentration of an RBC is extracted from the mean RI value of the cell. Because RBCs are primarily composed of Hb solution, the Hb concentration can be calculated using the linear relationship between the RI of Hb solution and the Hb concentration. Consequently, the cellular Hb content can also be obtained, since the cell volume and Hb concentration are provided. The dynamic membrane fluctuation is related to the deformability of an RBC. The dynamic membrane fluctuation is obtained using the mean RI and the temporal phase deviation, because the optical phase delay is identical to the integration of the RI and the thickness. A flow diagram showing the extraction of the properties is given in Fig. 1E. An example of single-RBC profiling is presented in Fig. 1F.

### 2.2. RBCs under different pathophysiological conditions

We acquired QPI images of RBCs drawn from individuals under well-defined pathophysiological conditions. The conditions included a healthy condition, IDA, RET, HS, and DM. IDA, which is one of the most prevalent diseases globally, is caused by insufficient dietary iron absorption compared to the iron demand of the body (Killip et al. 2007). RET is a syndrome related to failures in maintaining the functional RBC population, as reticulocytes are immature RBCs (Koepke and Koepke 1986). Pathological conditions including hypersplenism (Kar et al. 1986) and hemolytic anemia (Watanabe et al. 1994) have previously been related to an increase of reticulocytes. HS is an inherited disorder, characterized by the presence of spherical RBCs, or spherocytes, which are known to arise from defects in the linkages between the skeleton and the lipid bilayer of the RBC membrane (Perrotta et al. 2008). Clinically, HS is known to be related to splenomegaly, jaundice, and other hemolytic diseases. DM is a group of diseases characterized by hyperglycemia, originating in defects in insulin secretion or insulin functionality (Association 2010). DM is conventionally diagnosed or monitored according to the fasting plasma glucose level and glycated Hb level (Droumaguet et al. 2006).

The single RBC parameters measured using QPI are shown in Fig. 2. A total of 1028 RBCs were profiled using cDOT. The numbers of single-RBC profiling data items are 437, 99, 166, 86, and 240 for the healthy condition, IDA, RET, HS, and DM, respectively. The numbers of examined individuals are 7, 1, 1, 1, and 6 for the pathophysiological conditions in the same order.

**Fig. 2.**
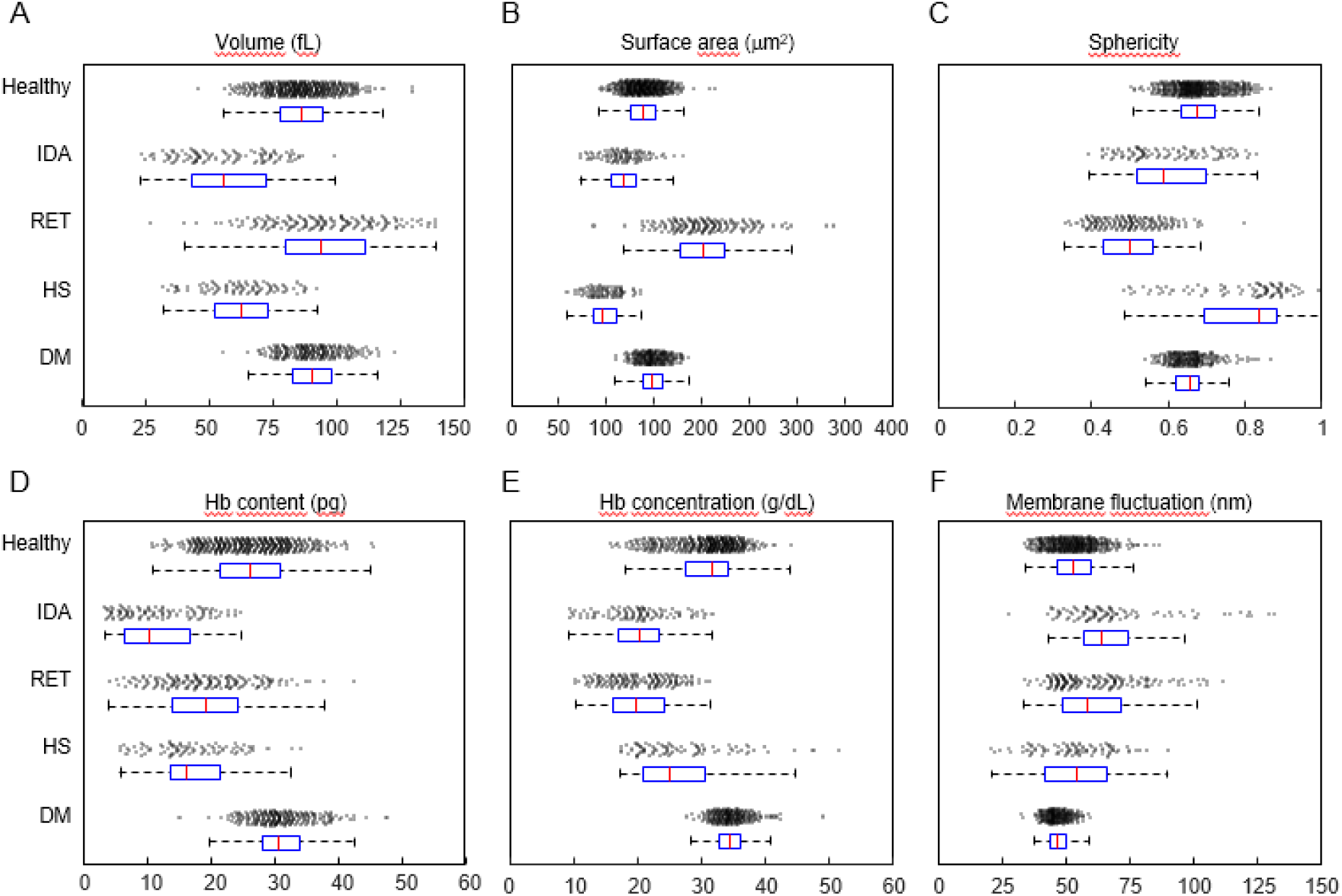
Properties of red blood cells obtained using single-cell profiling. (A) The volume, (B) surface area, (C) sphericity, (D) Hemoglobin (Hb) content, (E) Hb concentration, and (F) membrane fluctuation of red blood cells (RBCs) are plotted by the pathophysiological condition. IDA, RET, HS, and DM refer to iron-deficiency anemia, reticulocytosis, hereditary spherocytosis, and diabetes mellitus, respectively. Each population is described using a scatterplot and a box plot. The blue box of each distribution corresponds to the interquartile range (IQR) of that distribution. The red line in each box represents the median of the distribution. Each whisker is extended to the extremum unless the length is 1.5 times the length of the IQR.

This study and all experimental protocols were approved by the ASAN Medical Center Institutional Review Board (IRB project number: #IRB-13-90) and the KAIST Institutional Review Board (IRB project number: 2012-0128). Blood samples were collected over the regular course of patient care in the ASAN Medical Center, and we selected patients who had provided written informed consent for using their archival tissues for genetic testing. All data was de-identified.

The single-RBC profiling data was divided into a training set and a test set, to demonstrate disease screening based on unknown inputs. We randomly designated 40 profiling data items of RBCs under each pathophysiological condition as the test set, and employed the remaining data as the training set. Furthermore, the imbalance in sizes for the pathophysiological conditions in the training set was remedied using the synthetic minority over-sampling technique, in which the minority data is oversampled using linear combinations of the nearest neighbors (Chawla et al. 2002). The resulting size of the training set was 397 for each pathophysiological condition.

### 2.3. Neural network for predicting the pathophysiological condition based on single-RBC profiling

We implemented neural networks to screen for the pathophysiological conditions from input features consisting of the six RBC properties. The architecture of the neural networks is visualized in Fig. 3. A neural network classifier consisted of three fully-connected hidden layers of size 32, with rectified linear units (ReLUs) as the activation functions and a fully-connected output layer of size five with softmax activation, which normalizes the neural activation into probabilities. In order to accelerate the training and reduce the internal covariate shift, batch normalization was performed before each ReLU function (Ioffe and Szegedy 2015). The classifier was implemented using the TensorFlow 1.0.1 library for Python 3.5.3. The training and tests were run on a single central processing unit (Core i5-4670, Intel, USA).

**Fig. 3.**
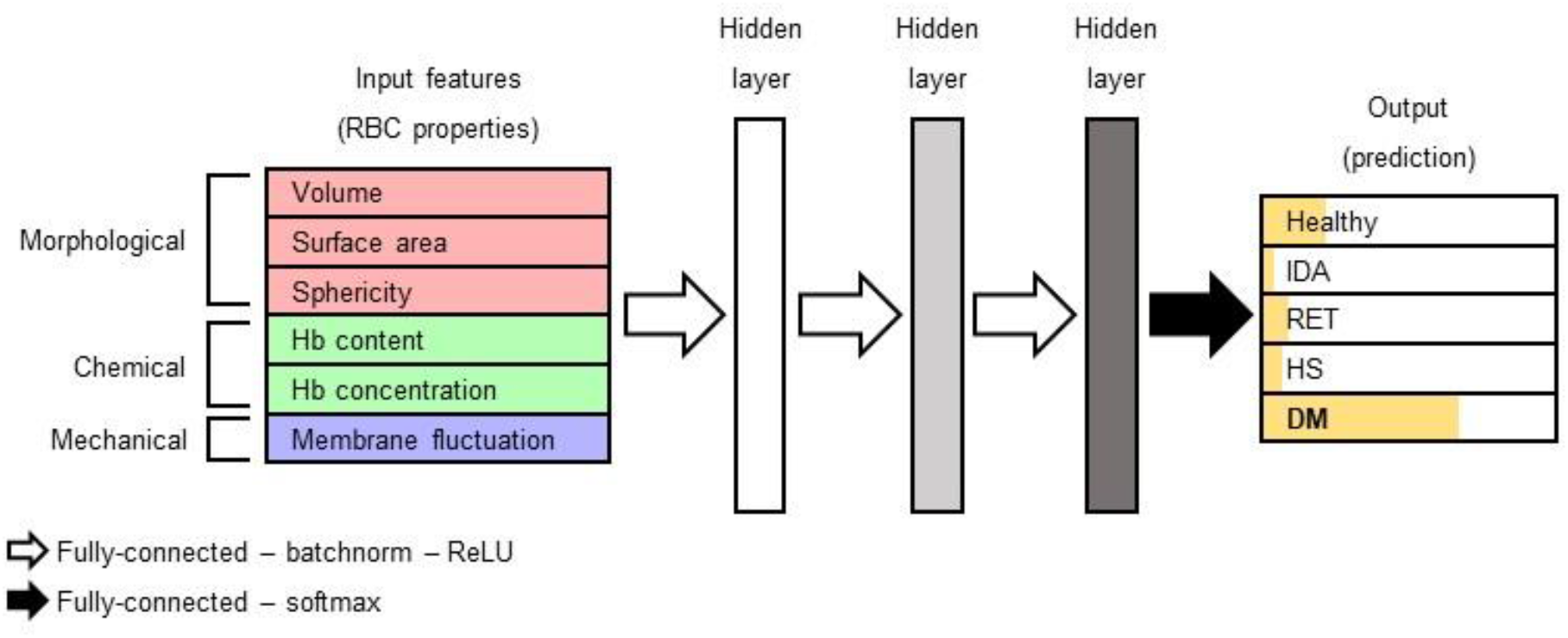
The architecture of neural networks for disease screening based on the profiling of single red blood cells. The neural network classifier predicts the pathophysiological conditions of the donor of a red blood cell (RBC) using the cellular properties of the RBC. The properties of the RBC are fed into the neural network as the input vector. Three hidden layers, each consisting of 32 elements, process the input vector. The output vector represents the probability of each physiological condition. The neural nodes are fully connected between each adjacent layer. Batch normalization and rectified linear unit (ReLU) activation follow the neural connection leading to each hidden layer, whereas softmax activation follows the neural connection leading to the output vector.

The neural networks were trained using the single-RBC profiling data of the training set. Before training, each element of each input vector was normalized such that the average and standard deviation for the training set became 0 and 1, respectively. During training, a loss function was minimized using the Adam optimizer (Kingma and Ba 2014), with a learning rate of 0.0001. The loss function was composed of a cross-entropy loss term and an L2 regularization term, in order to optimize the network for the training set and reduce over-fitting, respectively. A neural network was trained for 200 epochs, taking approximately an hour.

During the test, each single-RBC profiling data item of the test set was classified as one of the pathophysiological conditions. Based on the neural network output, which represents the probabilities of the five conditions, the most likely condition was chosen as the prediction result. The performance of the neural network classifier was verified to be higher than five other types of classifier (Fig. s1).

The prediction result for the test set was analyzed focusing on two aspects: the performance and the contributions of different input features. We evaluated the performance of the classifier using the sensitivity, specificity, and confusion matrix of the prediction for the test set (Table 1). Furthermore, we conducted an ablation study of the input features to identify the relevance of different types of RBC properties to screening for diseases and syndromes (Fig. 4).

**Fig. 4.**
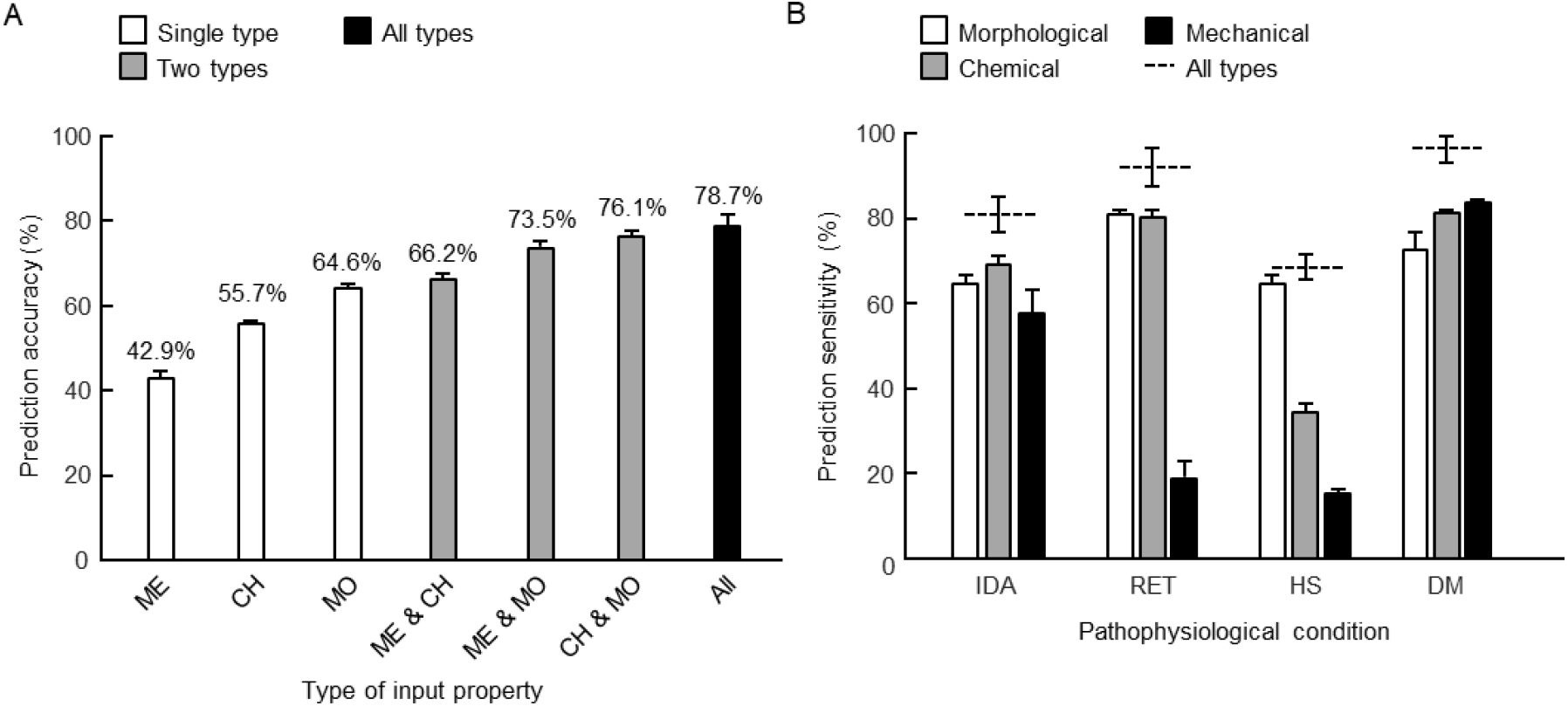
Prediction of pathophysiological conditions using different types of red blood cell properties. (A) The prediction accuracies plotted according to the types of red blood cell (RBC) properties that are used to train the neural network classifier. ME, CH, and MO refer to the mechanical, chemical, and morphological properties of RBCs, respectively. (B) The prediction sensitivities of pathophysiological conditions when the classifier is trained using single categories of the RBC properties. The plots are displayed in the form of the average ± standard deviation.

**Table 1.** Pathophysiological conditions predicted based on the profiling of single red blood cells.

|  | Prediction |  |  |  |  | Sensitivity |
| --- | --- | --- | --- | --- | --- | --- |
| Ground truth | Healthy | IDA | RET | HS | DM |  |
| Healthy | 23 | 1 | 1 | 1 | 14 | 57.5% |
| IDA | 5 | 33 | 0 | 2 | 0 | 82.5% |
| RET | 2 | 0 | 38 | 0 | 0 | 95.0% |
| HS | 0 | 12 | 0 | 28 | 0 | 70.0% |
| DM | 2 | 0 | 0 | 0 | 38 | 95.0% |
| Specificity | 94.4% | 91.9% | 99.4% | 98.1% | 91.3% | 80.0%<br>(Accuracy) |

### 2.4. Utilizing profiling data of multiple RBCs for the prediction of pathophysiological conditions

For accurate screening of diseases from single cells, it is necessary to overcome the cellular variation within an individual. While single-cell profiling based on QPI data provides an accurate method for characterizing individual cells, the distribution of RBC properties under an identical pathophysiological condition is broad. Incorrect predictions may occur for RBCs with properties that are dissimilar from the rest of the relevant population.

To overcome and exploit the wide spectrum of RBC populations, the classifier outputs of multiple RBCs were integrated for each individual. We attempted to increase the accuracy when examining an individual by statistically suppressing errors of predictions based on single-RBC profiling, while maintaining the utilization of single-RBC profiling. Specifically, the output vectors of the neural network were averaged over multiple RBCs belonging to the test set of an individual. The combinations of multiple RBCs were randomly chosen without replacement. This process is illustrated in Fig. 5, which presents an example of screening for DM with classification results of two RBCs. The improvement in the prediction performance obtained using the integration of multiple outputs is illustrated in Fig. 6. The performance using outputs of 10 RBCs is evaluated in detail using the sensitivity, specificity, and confusion matrix of the prediction for the test set (Table 2).

**Fig. 5.**
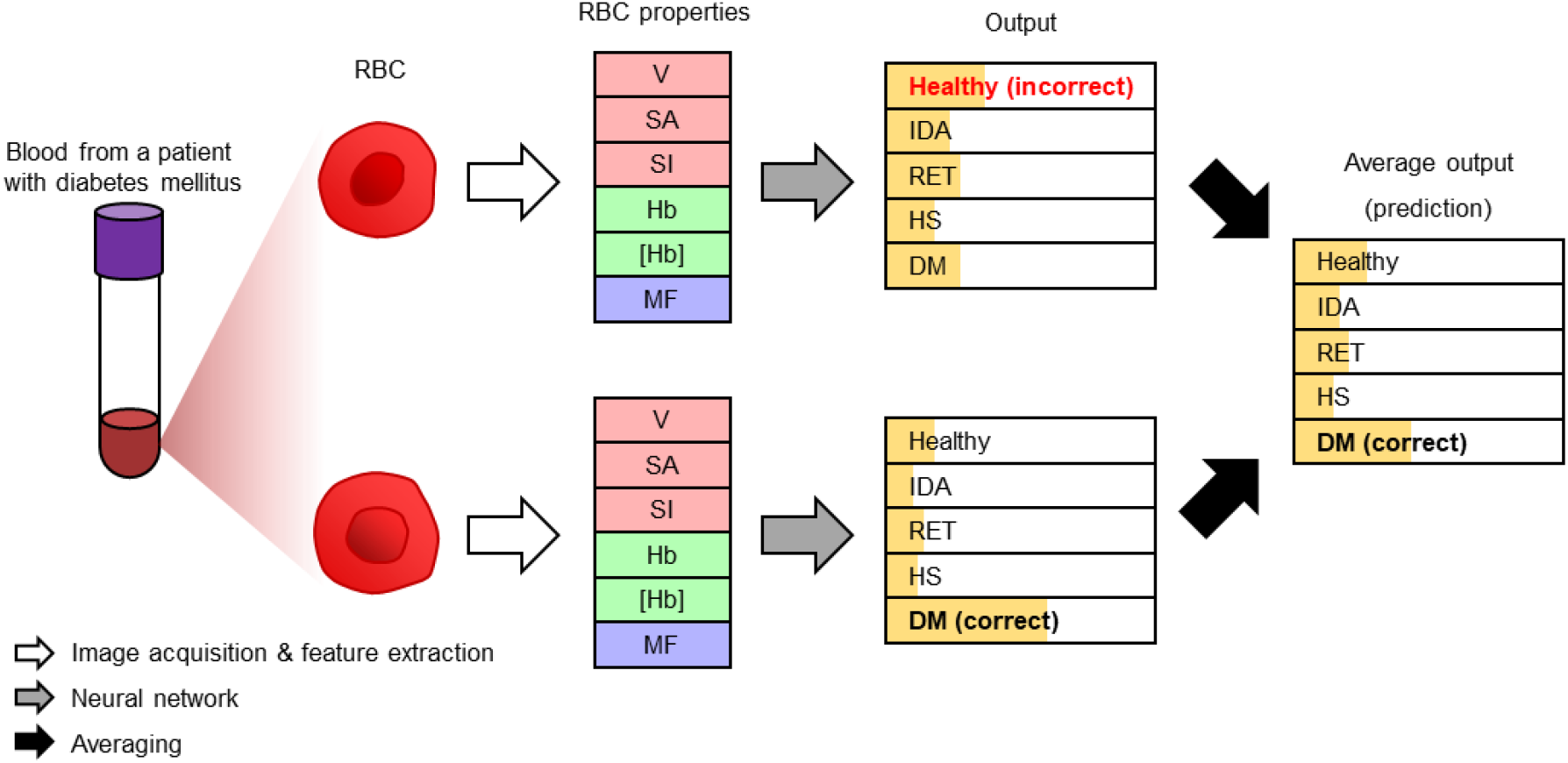
Schematic diagram of utilization of profiling data for multiple red blood cells. The outputs for multiple red blood cells (RBCs) of an identical donor are averaged over. The diagram displays an example of screening for DM in the presence of an incorrectly classified RBC. V, SA, SI, Hb, [Hb], and MF refer to the volume, surface area, sphericity, hemoglobin (Hb) content, Hb concentration, and membrane fluctuation, respectively.

**Table 2.** Pathophysiological conditions predicted based on the profiling of 10 red blood cells.

|  | Prediction |  |  |  |  | Sensitivity |
| --- | --- | --- | --- | --- | --- | --- |
| Ground truth | Healthy | IDA | RET | HS | DM |  |
| Healthy | 38 | 0 | 0 | 0 | 2 | 95.0% |
| IDA | 0 | 40 | 0 | 0 | 0 | 100% |
| RET | 0 | 0 | 40 | 0 | 0 | 100% |
| HS | 0 | 0 | 0 | 40 | 0 | 100% |
| DM | 0 | 0 | 0 | 0 | 40 | 100% |
| Specificity | 100% | 100% | 100% | 100% | 98.8% | 99.0%<br>(Accuracy) |

## 3. Results and discussion

### 3.1. Properties of RBCs under pathophysiological conditions

The six cellular properties of each RBC were extracted from the 3D RI tomogram and time-lapse phase maps. The volume, surface area, sphericity, Hb content, Hb concentration, and membrane fluctuation are plotted for the RBCs according to the pathophysiological condition in Fig. 2. The average ± standard deviation volumes of the RBCs were 88.1 ± 13.0, 56.3 ± 17.7, 94.6 ± 21.5, 62.3 ± 14.5, and 90.2 ± 10.9 fL for the healthy individuals, IDA patient, RET patient, HS patient, and DM patients, respectively. In the identical order, the surface areas of the RBCs were 143.1 ± 17.9, 117.8 ± 21.2, 202.6 ± 38.2, 97.8 ± 16.7, and 148.2 ± 14.4 μm^2^; the sphericities of the RBCs were 0.67 ± 0.06, 0.60 ± 0.11, 0.50 ± 0.08, and 0.66 ± 0.05; the Hb contents were 27.3 ± 6.4, 11.7 ± 5.9, 19.4 ± 7.3, 17.1 ± 6.3, and 31.2 ± 4.8 pg; the Hb concentrations were 30.9 ± 4.6, 19.9 ± 5.3, 20.1 ± 5.0, 27.2 ± 7.5, and 34.6 ± 2.7 g/dL; and the membrane fluctuations were 51.0 ± 8.6, 68.9 ± 20.6, 61.6 ± 16.5, 53.5 ± 14.4, and 46.6 ± 4.5 nm.

It was evident from the distributions of the RBC properties extracted from the QPI images that a tool for high-dimensional analysis is essential for the classification of RBCs. While the population-average of a property varies between the pathophysiological conditions, the range of the entire population significantly overlapped between the conditions. The pathophysiological condition of an RBC was unlikely to be accurately predicted when only a single property of the RBC was provided.

### 3.2. Screening for diseases and syndromes based on profiling single RBCs

The trained neural networks were capable of predicting the pathophysiological condition from the properties of single RBCs for the majority of the investigated RBCs. The average test accuracy of the classification was 78.7% over five neural network models that were trained independently. Table 1 presents the prediction results for the test set obtained using one of the five trained models. Here, each row represents the ground truth condition and each column represent the predicted condition. The diagonal elements refer to correct predictions, whereas the off-diagonal elements refer to incorrect predictions. The neural network classifier was relatively more sensitive to RET and DM, for which the sensitivities of screening were over 90%, compared with other pathophysiological conditions. The most frequent error was the misclassification of RBCs of healthy individuals as RBCs of DM patients, which is not a failure to screen for unhealthy conditions, but a case of false alarm. Another case of error was misclassifying the RBCs of the HS patient as the RBCs of IDA patients.

The prediction process for an unknown input vector was rapid using a trained neural network, taking less than one second. Such rapid inference is the advantage of using an eager learning algorithm such as a neural network. In contrast to a lazy learning algorithm, for which the target function must be locally approximated for each input, the target function of an eager learning algorithm is approximated during training, which excludes the necessity of heavy computations. We consider this as an advantage for disease screening tasks.

### 3.3. Ablation study of the input features

The contributions of different types of RBC properties to the screening of the pathophysiological conditions was investigated by conducting an ablation study of the input features. The performances of the neural network classifiers trained using different types of input features were evaluated. The input features were classified into three types: morphological, chemical, and mechanical properties of RBCs. A total of seven cases were compared, comprising three cases of a single-type input, three cases of a two-type input, and the entire input case. The prediction accuracies for the test set in the above-mentioned cases are displayed in Fig. 4A. The accuracies were 42.9, 55.7, 64.6, 66.2, 73.5, 76.1, and 78.7% when using the mechanical properties, chemical properties, morphological properties, mechanical and chemical properties, mechanical and morphological properties, chemical and morphological properties, and all three properties, respectively.

The results of the ablation study indicated that all three types of RBC properties are involved in screening for the pathophysiological conditions. The prediction accuracy increased as we utilized more types of RBC properties for the training, while it did not increase in any case of excluding a type of feature. This reflects the fact that no single type of RBC property can be fully derived from the other two, owing to the fundamental differences between the three types of RBC properties.

Furthermore, the three types of RBC properties did not contribute equally to the screening for the pathophysiological conditions. In the three cases using single categories of RBC properties, the accuracies of the prediction were significantly different. The morphological properties appeared to be the most critical features in screening for the investigated conditions, because the presence of morphological properties in the input vector increased the classification accuracy more than the other two types of RBC properties. The chemical features were indicated to be the next most important features, based on the identical criterion. However, the presented order of importance does not determine the universal priorities for representing RBCs from the pathophysiological perspective. The magnitudes may vary depending on the investigated conditions, because different diseases and syndromes involve different alterations in RBCs.

Furthermore, the key traits of the RBCs that the neural network selects to screen for each unhealthy condition appear to be consistent with the literature. We studied the role of each RBC property type in the screening for the unhealthy conditions by examining the disease sensitivities of neural networks trained using single categories of RBC properties (Fig. 4B). Each condition was most sensitively screened for by the neural network trained using a specific category of RBC properties, even though the morphological properties were the most critical to the overall prediction accuracy. IDA, RET, HS, and DM were most sensitively screened for when the classifier was trained using the chemical, morphological, morphological, and mechanical properties, respectively. The aforementioned result is in agreement with the previous findings that IDA reduces the average Hb contents of RBCs (Killip et al. 2007), RET increases the average volumes of RBCs (Koepke and Koepke 1986; Watanabe et al. 1994), HS increases the average sphericities of RBCs (Kim et al. 2014d; Perrotta et al. 2008), and DM suppresses the average membrane fluctuations of RBCs (Lee et al. 2017; Schmid-Schönbein and Volger 1976).

### 3.4. Screening for diseases and syndromes based on profiling multiple RBCs

The accuracy of predicting pathophysiological conditions of individuals was further increased by utilizing the neural network outputs of multiple RBCs. The output vector was averaged over multiple RBCs of an individual to predict the condition. The accuracies when averaging the output vectors between 1 and 10 RBCs are presented in Fig. 6A. The overall accuracy increased from 78.7 ± 2.5% to 98.8 ± 0.3%. The slope of the increase in the prediction accuracy was steeper for smaller numbers of RBCs. Fig. 6B and C display the confusion matrices of the predictions based on single RBCs and averages of 10 RBCs respectively. Comparing the two 3D bar graphs shows a reduction in the number of incorrect predictions when averaging over the output vectors of 10 RBCs, as off-diagonal elements in the graph are conspicuously removed. Table 2 specifies the detailed prediction results using the output averaging method, where the overall accuracy was 99.0%. Here, the only remaining errors comprised misclassifications of healthy individuals as DM patients, which is in line with the most frequent errors in the prediction using single RBCs.

**Fig. 6.**
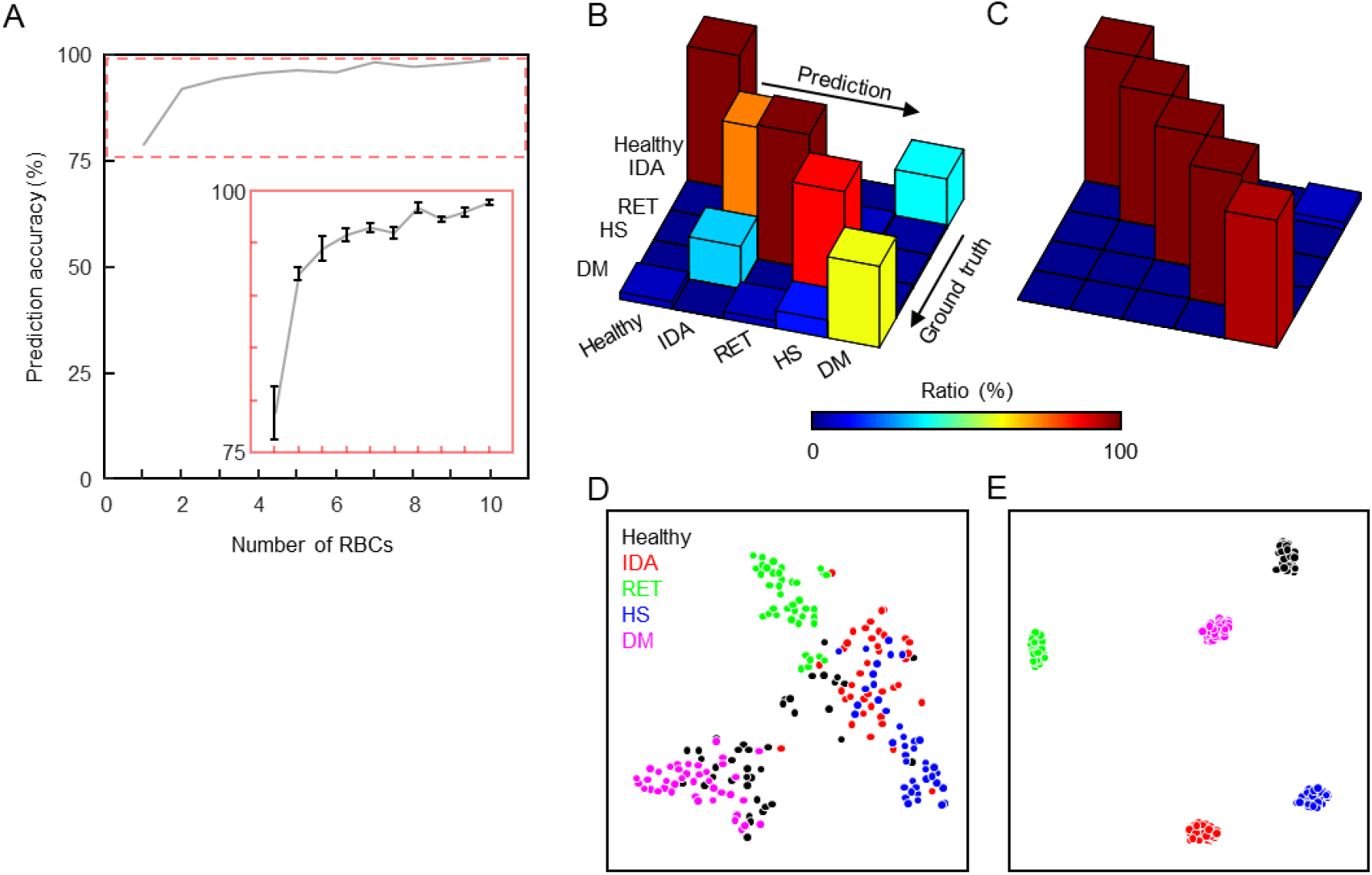
Influence of utilizing multiple outputs of the neural network. (A) The accuracy of predicting the pathophysiological conditions according to the number of red blood cells (RBCs) over which the classifier outputs are averaged. The inset shows the rescaled plot of the range displayed using the dashed line. (B) Visualization of the confusion matrix resulting from the prediction based on the profiling of single RBCs. (C) Visualization of the confusion matrix resulting from the prediction based on the profiling of 10 RBCs. (D) T-distributed stochastic neighbor embedding (t-SNE) visualization of the activation at the final hidden layer. (E) T-SNE visualization of the 10-fold averaged activation at the final hidden layer.

The increase in the accuracy indicates that the average neural activations converge to within a narrower scope, even though the neural activations of individual RBCs lie in a wide space. This indication was underpinned by visualizing the distributions of neural activations in the final hidden layer using t-distribution stochastic neighbor embedding (van der Maaten and Hinton 2008) (Fig. 6D and E). While the activations of the final hidden layer are not fully separated according to the pathophysiological condition, the average values of the activations are well separated.

## 4. Conclusion

We have demonstrated accurate screening for diseases and syndromes by combining QPI-based single-RBC profiling with machine learning. Our data-driven investigation of the high-dimensional feature space of a single-cell profiling dataset resulted in a high-accuracy neural network classifier, which systematically integrates morphological, chemical, and mechanical properties of individual RBCs. The pathophysiological condition-specific fingerprint patterns learned by the classifier were consistent with the literature. Combining the network predictions for multiple RBCs led to an accuracy of over 98% by simply averaging over 10 RBCs, implying the fruitfulness of information available from QPI.

The proposed method exploits the inherent advantages of both QPI and machine learning. First, the label-free nature of QPI minimizes the required sample preparation, and thus enables rapid single-cell profiling. Second, QPI-based profiling of individual RBCs provides unique information that is otherwise inaccessible (i.e., membrane fluctuation, which is itself an important research topic in biophysics but has not been utilized in previous screening methods). Third, when trained beforehand a trained neural network classifier can perform extremely fast predictions immediately following the QPI measurement. Altogether, rapid and efficient screening for multiple diseases and syndromes was achieved.

The present work has several limitations, which can be improved in future work. Above all, it is essential to combine the QPI system with high-throughput microfluidics for generating large-scale datasets in order to explore the questions arising from the current work. For example: How many pathophysiological conditions can be screened? Is it possible to screen for multiple diseases and syndromes simultaneously? How many RBCs are required for good enough screening? Is there a systematic strategy to explore the learned patterns in the feature space for a data-driven understanding of diseases and syndromes? Furthermore, such large-scale data would enable an advanced implementation of deep neural networks or deep learning (Jo et al. 2017), with a stronger learning ability provided through feature learning. While the proposed machine learning framework is based on the hand-crafted features from previous studies on RBCs, deep learning automatically learns and extracts more powerful features from raw images. Furthermore, using commercially available QPI techniques we expect that the present method can be readily applied to clinical medicine (Shin et al. 2016). Finally, we expect that cost-effective miniaturization of the system would enable rapid point-of-care diagnosis.

## Acknowledgment

The authors thank Dr. Youngchan Kim (National Institutes of Health) and Mr. SangYun Lee (KAIST) for providing the cDOT dataset. This work was supported by BK21+ program, and National Research Foundation of Korea [2015R1A3A2066550, 2017M3C1A3013923, 2014K1A3A1A09063027]. Y. Jo acknowledges support from KAIST Presidential Fellowship and Asan Foundation Biomedical Science Scholarship.

